# Quality-quantity tradeoffs drive functional trait evolution in a model microalgal “climate change winner”

**DOI:** 10.1101/819326

**Authors:** Rasmus T. Lindberg, Sinéad Collins

## Abstract

Phytoplankton are the unicellular photosynthetic microbes that form the base of aquatic ecosystems, and their responses to global change will impact everything from food web dynamics to global nutrient cycles. Some taxa respond to environmental change by increasing population growth rates in the short-term, and, based on this, are projected to increase in frequency over decades. To gain insight into how functional traits in these projected “climate change winners” change over different timescales, we evolved populations of microalgae in ameliorated environments for several hundred generations. While populations initially responded to environmental amelioration by increasing photosynthesis and population growth rates as expected, this response was not sustained. Instead, most populations evolved to allocate a smaller proportion of carbon to growth while increasing their ability to tolerate and metabolise reactive oxygen species (ROS). This diversion of fixed carbon from growth to catabolism underlies a quality-quantity tradeoff in daughter cell production which drives the evolution of population growth rates and of functional traits that underlie the ecological and biogeochemical roles of phytoplankton. There is intraspecific variation in the trait combinations that evolve, but all are consistent with mitigating ROS production and accumulation in ameliorated environments over hundreds of generations. This offers both an evolutionary and a metabolic framework for understanding how functional traits can change in primary producers projected to be “climate change winners”, and suggests that short-term population booms and associated trait shifts have the potential to be dampened or reversed if environmental amelioration persists.

---

Microbial primary producers (phytoplankton) form the base of aquatic food webs and fix approximately 40% of global carbon annually (1). Photosynthetic microbes respond to increases in carbon dioxide using plastic responses (short term change in phenotype with no underlying change in genotype) (2, 3) and evolutionary responses (change in the genetic composition of a population over time) (4, 5). Phytoplankton can be grouped based on their roles in biogeochemical cycles (calcifyers, silicifyers, nitrogen fixers, other green), and there is variation in both plastic and evolutionary responses to increased CO_2_ availability within and between phytoplankton functional groups (2, 3, 6, 7). This variation has led to the prediction that taxa that can respond to nutrient enrichment (such as increased carbon availability) by increasing their population growth rate will increase in frequency as these conditions persist (2, 8). These taxa are often called “winners” under global change, and it is expected that their influence on food webs and nutrient cycles grow as winners increase in frequency (9), and as eutrophication increases (10)(11). While predictions about the composition and function of future microbial communities of primary producers are based on the increasing importance of lineages that subjectively experience an environmental change as improvement, our understanding of evolutionary processes is based predominantly on experiments carried out in stressful, starvation, or toxic environments (12). A nuanced understanding of within-lineage evolution under many aspects of global change thus requires investigating how natural selection acts when organisms initially experience an environmental change as an improvement over their historical environment.

When short-term trait changes are used to project the characters of functional groups over longer timescales (years or decades), it is assumed that plastic increases in lineage growth rate, alongside changes in functional traits such as photosynthesis, respiration, and cell size, that occur in the enriched environment can be sustained indefinitely. However, theoretical (13, 14) and limited empirical (15–18) support for this assumption is equivocal (but see (19)). We reason that when populations are exposed to permanently ameliorated environments, they have probably previously experienced those environmental improvements, or similar ones, episodically. For environmental ameliorations such as increases in pCO_2_ or nutrient enrichment, cells can express a plastic response to the new environment that appears (or is known to be) adaptive, given that the environmental improvement is transient. For example, while increasing metabolic and thus cell division rates over a few generations can be adaptive, maintaining high metabolic rates is also associated with the increased production of toxic metabolic byproducts, such as reactive oxygen species (ROS) (20) that can damage cellular components including proteins, nucleic acids and membranes (21). Limiting ROS damage is thought to play a role major evolutionary transitions such as the evolution of dedicated germ lines and asymmetric cell division, and can limit the upper rate of cell division possible in organisms without a dedicated germ line (13, 22–24). Similarly, we have shown using simulations that the accumulation of toxic metabolic byproducts can limit population growth rates of unicells in ameliorated environments, and lead to reductions in population growth rate if reallocating energy from growth to repair increases daughter cell survival. This is called Prodigal Son dynamics – an adaptive decrease in population growth rate as a consequence of a previous sustained increase in cell division rates (14). Based on this, we predict that selection in ameliorated environments eventually shifts from short term (plastic) responses based on producing many daughter cells to long-term (evolutionary) responses that produce higher quality (less damaged) daughter cells.

To investigate how lineage growth rate evolves in ameliorated environments, we evolved five genotypes of the green alga *Chlamydomonas reinhardtii* in single genotype cultures for over 900 generations in nutrient-rich, high pCO_2_ environments. Evolved populations from the high pCO_2_ environment are called “High” (for High-evolved) populations, and those evolved in the ambient pCO_2_ environment are called “Amb” (for Ambient-evolved) populations. Our previous empirical (15–17, 25, 26) and modelling (14) studies showed that decreases in population growth rates could occur under these conditions, but did not provide a general mechanism driving trait reversion in ameliorated environments. Here, we look for evidence of a quality-quantity tradeoff in daughter cell production by measuring or calculating several traits associated with life history (daughter cell production, cell cycle length), daughter cell quality (ROS tolerance and heat shock recovery). We show that this shift from producing many to higher-quality daughter cells is associated with shift in traits that are commonly used to understand the role of microbial primary producers in aquatic ecosystems (photosynthesis and respiration rates, cell size, biovolume production).

## Results and Discussion

### Plastic responses to environmental enrichment

Because we were interested in the evolutionary consequences of sustained rapid growth exceeding that in a benign, nutrient-replete environment that populations were historically grown in, we used 2000ppm pCO_2_ and continuous light in nutrient-replete media as our ameliorated environment. This results in a population growth rate increase of ~12% relative to the same media and conditions with ambient pCO_2_, which is near the maximum reported for this species (27). These are not intended to be realistic environments, but rather ones that represent a standard, non-stressful control or ancestral environment (nutrient replete ambient pCO_2_) and an improved environment that allows a substantial increase in population growth rate (nutrient replete high pCO_2_). Further increases in nutrients, light and pCO_2_, or changes in general culturing conditions, did not further increase population growth rates.

Plastic responses to environmental enrichment were measured by growing populations in the nutrient-rich, high pCO_2_ environment for two transfers (~9 cell divisions). Plastic responses can be seen in Fig 1 by comparing the Amb populations growing in ambient pCO_2_ (pink, left hand boxes of each plot) with the Amb populations growing in high pCO_2_ (green, middle boxes of each plot). As expected, population growth rate and gross photosynthesis rates increase in the enriched environment in agreement with other studies on this and other microbial systems in marine and freshwater systems (2, 3, 7, 17)(effect of pCO_2_ on population growth rate t_54_=8.18, p <0.0001 and gross photosynthesis rate per cell t_54_=9.54, p <0.0001). Plastic responses in respiration also occur in most genotypes, though the direction of response varies between genotypes. In phytoplankton, growth depends crucially on two metabolic fluxes, gross photosynthesis (P, the gross fixation of inorganic carbon (C)) and respiration (R, the gross remineralization of organic C) change(28); carbon use efficiency (CUE = 1 − R/P), is the fraction of C available for growth after the catabolic demands of the cell have been met. Carbon use efficiency, and thus the amount of C being allocated to growth relative to non-growth in cells, increases during plastic responses to increased pCO_2_ in 4/5 genotypes. The single genotype (cc1691) that slightly decreases CUE under CO_2_ enrichment has high CUE under both ambient and high pCO_2_ relative to other genotypes. Further increasing CUE may not be possible in this experiment, since no genotype exceeds these CUE values under any conditions.

**Figure 1.**
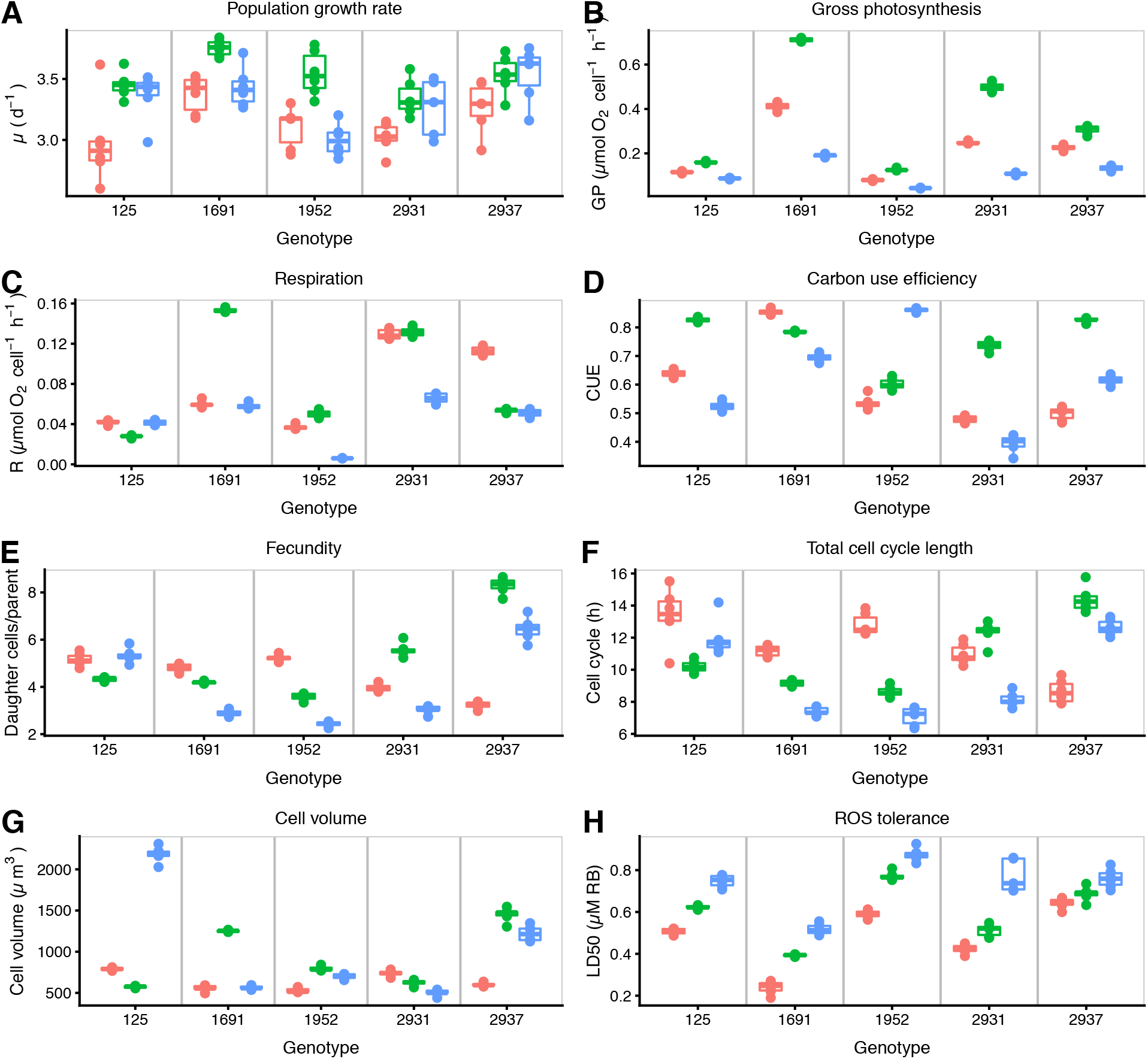
Control, plastic, and evolved trait values for functional traits. Trait values in pink correspond to the Amb populations growing in ambient pCO_2_, and correspond to the baseline trait prior to environmental amelioration. Trait values in green show Amb populations growing in high pCO_2_, which corresponds to the plastic response to environmental amelioration. Trait values in blue show High populations growing in high pCO_2_, and are the evolved phenotype after ~900 generations in the ameliorated environment. Each point represents the average trait value for a single replicate population, boxes show means for each genotype.

To understand the underlying life history strategies that led to plastic increases in overall population growth rate, we also measured the number of daughter cells produced per cell cycle, calculated the time needed for one complete cell cycle, and measured shifts in average cell size (Fig 1 panels EG). Interestingly, both the strategy of increasing total cell cycle length and producing more daughter cells (increasing fecundity but reproducing less often) and decreasing total cell cycle length but producing fewer daughter cells (decreasing fecundity but reproducing more often) occur in response to CO_2_ enrichment in *C.reinhardtii*, and strategies are genotype-specific. Both strategies increase population growth growth rate (μ). While cell size also responds to environmental enrichment, changes are not associated with any particular strategy or rate of population growth increase. Previous work with single genotypes of *C.reinhardtii* has found that the number of daughter cells produced per cell cycle was positively correlated with parent cell size within single populations of that genotype (29). In our study, differences in average cell size do not explain differences in the number of daughter cells produced between different genotypes.

### Evolutionary responses to environmental enrichment

All High populations were grown in the high or ambient pCO_2_ environments for 90 transfers, corresponding to ~900 generations in the High populations. Evolved trait values are shown as the trait values of the High populations grown at high pCO_2_ (Fig 1, blue, right-hand boxes of each plot), so are a combination of evolutionary and sustained plastic responses.

By the end of the experiment, High populations had, on average, decreased their population growth rate relative to Amb populations (effect of selection environment t_82_= −4.11, p = 0,0001, assay environment t_82_ = 2.02, p = 0.046, and interaction term t_82_ = −2.28, p = 0.025, on population growth rate). This is driven by the two genotypes that reproducibly reduced their growth rate at high pCO_2_ relative to Amb populations. See Fig 1A. These data show that environmental amelioration by CO_2_ enrichment can drive evolutionary responses, in line with previous studies in this and other microalgal systems (17, 19, 30, 31). There is intraspecific (between genotype) variation in both plastic and evolutionary responses to CO_2_ enrichment in *C. reinhardtii*. Despite this variation, there is a pattern where the evolutionary responses can decrease, but not increase, population growth rates relative to the plastic trait value, such that High populations never grow faster, and sometimes grow more slowly than, Amb populations of the same genotype in high pCO_2_. This partial or complete reversal of the plastic growth response to CO_2_ enrichment is consistent with decreased population growth rates being part of an adaptive phenotype in the ameliorated environment when it occurs, presumably because growth is correlated with other traits that determine cell quality.

### Quality-quantity tradeoffs in cell production in enriched environments

Based on previous simulations (13, 14, 22, 23), we hypothesized that the number of daughter cells produced can be limited by their quality in the ameliorated environment. We further expected that quality be related to the ability to tolerate or detoxify metabolic byproducts such as reactive oxygen species (ROS). These data are summarized in Figure 1 and can be seen by comparing the plastic (green, middle bars) trait values with evolved (blue, right hand bars) trait values for each genotype and trait.

To test whether or not there was evidence of a quality-quantity tradeoff in daughter cell production driving decreased population growth rates in the High populations, we measured daughter cell production (cell fecundity) as the average number of cells produced per mother cell per cell cycle for the High and Amb populations (Figure 1E). Alongside the selection environment, the assay environment, and the interaction between those two (statistics above), the number of daughter cells modulates how the selection environment affects population growth rate (daughter cell × selection environment t_82_ = 4.00, p = 0.001) and the growth responses of evolved populations to changes in pCO_2_ (daughter cell × selection environment × assay environment t_82_ = −2.28, p = 0.025). In 4/5 genotypes, daughter cell production per cell cycle in high pCO_2_ decreased by ~1 cell. Regardless of whether the plastic response was to increase or decrease fecundity in the high pCO_2_, the evolutionary response decreased fecundity relative to the plastic trait value. In these 4 genotypes, total cell cycle length and average cell volume also decreased relative to the plastic trait values.

Overall, the evolutionary response to high pCO_2_ includes slowing down metabolism, as evidenced by lower gross photosynthesis and (for 4/5 genotypes) lower rates of respiration per cell relative to the plastic trait values (Fig 1B-C) (Effect of selection regime on gross photosynthesis rate per cell t_110_ = −2.52, p=0.013). Gross photosynthesis and respiration do not slow proportionally during evolution in the high CO_2_ environment. This is reflected by differences in the plastic and evolutionary CUE values (Fig 1D). Since photosynthesis unavoidably produces toxic metabolic byproducts such as ROS, and higher photosynthetic rates are often associated with higher ROS production (32), lower gross photosynthesis is consistent with decreasing photosystem II activity to limit ROS production. In 4/5 High genotypes, evolved CUE is also lower than the plastic trait value. This indicates that less C is being used for growth, and more for catabolism, in High populations than expected based on the plastic response. The proportion of C allocated to growth decreases over time under persistent environmental amelioration, even if the external environment is constant.

Along with reducing ROS production, a second possible way to respond to selection against ROS-induced damage would be to increase ROS tolerance or detoxification. We tested whether High populations more tolerant to exogenous ROS than Amb populations when both were grown in high pCO_2_, and found that they were (Figure 1H) (t_53_=11.07, p<0.001). We also found that recovery from heat shock of High populations when grown at high pCO_2_ was higher than recovery of Amb populations grown at high pCO_2_ (t_53_=8.35, p<0.001). The rationale for this measurement is that heat stress affects a wide range of cellular processes, such that low levels of damage that may have no phenotypic effect in benign environments may be revealed under heat stress (33). Together, these data show evidence of a quantity-quality tradeoff during evolution in an ameliorated environment. The data are consistent with the tradeoff being driven by selection against incurring damage from toxic metabolic byproducts, with ROS being one example of such a byproduct. Populations evolve to produce fewer, but higher-quality daughter cells per cell division cycle, than they do when they initially encounter the CO_2_ enriched environment. High populations also recovered more rapidly from heat-shock (see SI) indicating that daughter cell quality is not entirely environment-specific, as populations were never selected at high temperatures. Interestingly, short term exposure to high pCO_2_ increases ROS tolerance (Figure 1H) but decreases recovery from heat shock (SI Figure 1). This is consistent with ROS metabolism being stimulated, but cells being in worse overall condition or with energy being reallocated from other processes to ROS metabolism, even after only ~9 generations in high pCO_2_.

We investigated whether increased tolerance to exogenous ROS in high pCO_2_-evolved cells was consistent with upregulating antioxidant genes, which is a strong indication that the ability to detoxify ROS has increased. To do this, we measured gene expression levels for the production of key enzymes *Chlamydomonas* ROS metabolism in three of the five genotypes in this experiment (cc-1691, cc-2931 and cc-2937) for three genes involved in ROS metabolism, covering both the mitochondria and the chloroplast. Specifically, we measured the expression of *Cat*, *Fsd1* and *Gpx* genes, encoding Catalase, Iron-Superoxide Dismutase and Glutathione Peroxidase, respectively, in high pCO_2_. Gene products are located in either the mitochondrion (CAT), the chloroplast (FSD1), or both (GPX) (34–36). See Figure 2. While expression levels vary with genotype and gene product, we find that in general, High populations have upregulated expression of the ROS metabolism genes measured relative to the Amb populations of the same genotype when grown at high pCO_2_ (effects of selection on gene expression level *Cat*: t_14_=5.67, p=0.0001, *Fsd*1: t_14_=2.05, p=0.05, *Gpx*: t_14_=5.48, p=0.001). Previous studies have shown that upregulating *Gpx* alone is sufficient to enhance singlet oxygen resistance in *C.reinhardtii* (20). Overall, better heat shock recovery, higher tolerance to exogenous ROS and the upregulation of ROS metabolism genes is consistent with a lower CUE in High populations, where more carbon is being allocated to processes other than growth. This reallocation of energy from growth to an increased ability to mitigate damage is consistent with the driving mechanism of Prodigal Son dynamics proposed in our previous model (14).

**Figure 2.**
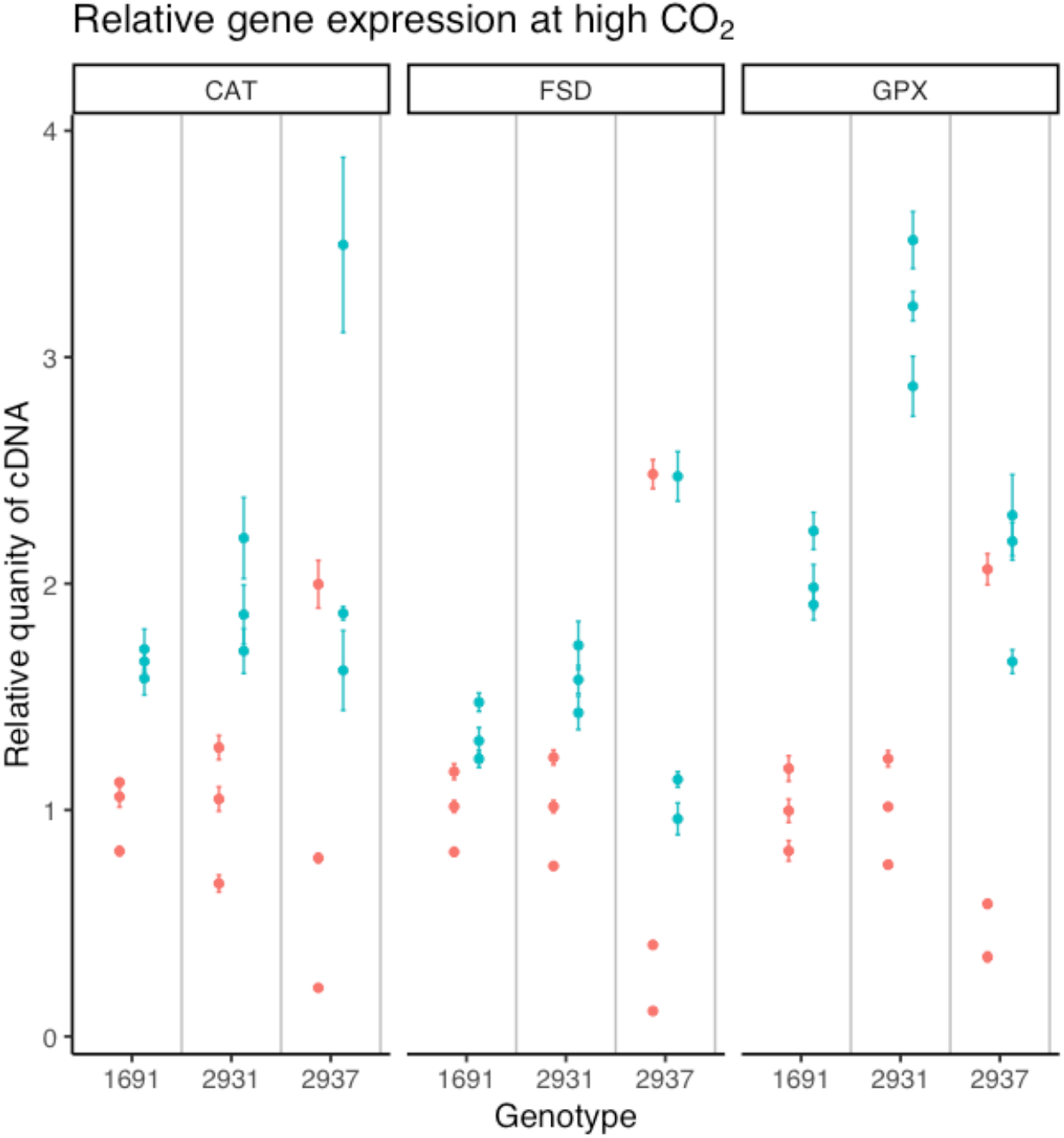
Gene expression levels of three antioxidant genes at high pCO_2_. CAT = Catalase, found in the mitochondria; FSD= Iron-Superoxide Dismutase, found in the chloroplast; GPX = Glutathione Peroxidase, found in both locations. Pink dots are for Amb populations growing at high pCO_2_, blue dots are for High populations growing at high pCO_2_. Each dot is a single independently evolved population, bars show SEM for independent replicate cultures and RNA extractions for each population. For each genotype and gene combination, the average value of the Amb population expression levels is set to 1.0; values shows are relative to this, and so are fold-change in expression level relative to Amb of the same genotype.

The majority of our experimental populations showed evidence of Prodigal Son dynamics when evolving in high pCO_2_, but there were exceptions. For example, cc125 High populations consistently maintain the same population growth rate and fecundity as the cc125 Amb populations in high pCO_2_, despite the High populations having lowered gross photosynthesis rates and higher ROS tolerance. The cc125 High populations also show different directions of trait evolution than the other four genotypes. This indicates that evolving higher ROS tolerance without a decrease in fecundity is possible.

### The impact of Prodigal Son dynamics on projecting functional traits of future populations

To understand how the evolved strategies for growing in a persistent high pCO_2_ environment can impact ecological function under environmental amelioration, we report our data as traits commonly used in aquatic ecosystem or biogeochemical models: cell size, biovolume (as a proxy for biomass) production per day, chlorophyll a production per day (37, 38). See Figures 1G and 3. For four of the five genotypes, cell volume is smaller in High populations than in Amb populations grown at high pCO_2_, which would affect both the sinking rate and grazibility of the cells, with potential effects on nutrient cycling and trophic interactions. In general, the total biovolume produced per day by High populations is lower than the Amb populations grown at high pCO_2_, which is consistent with most of the genotypes of High cells producing fewer, smaller daughter cells. One genotype, cc125, produces many large daughter cells. The total chlorophyll produced per day varies more over the 5 genotypes, illustrating that not only individual traits, but also trait correlations can evolve over time. For all traits, the range of trait values in the High populations is larger than the range in the Amb populations, suggesting that for modelling efforts, parameter value ranges based on plastic responses will underestimate the possible range of functional traits over time periods where adaptive evolution can also occur in ameliorated environments.

**Figure 3.**
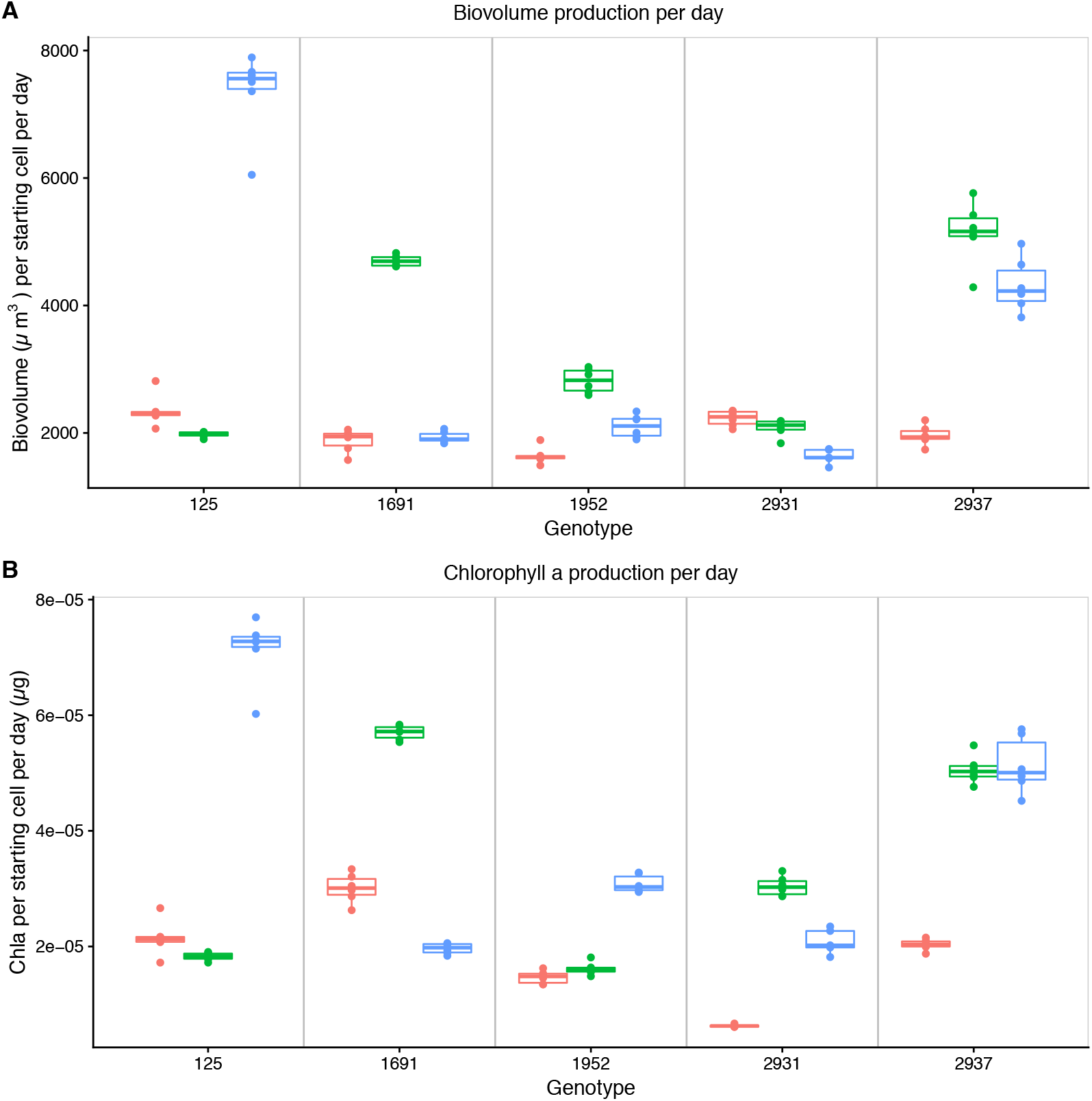
Measures of two proxies of primary production. Trait values in pink correspond to the Amb populations growing in ambient pCO_2_, and correspond to the baseline trait prior to environmental amelioration. Trait values in green show Amb populations growing in high pCO_2_, which corresponds to the plastic response to environmental amelioration. Trait values in blue show High populations growing in high pCO_2_, and are the evolved phenotype after ~900 generations in the ameliorated environment. Each point represents the average trait value for a single replicate population, boxes show means for each genotype. For both traits, values are calculated as the amount of either biovolume (A) or chlorophyll a (B) that would be produced starting from a single ancestral cell, through growth and cell division, over a 24 hour period.

Many aspects of global change are detrimental to large numbers of organisms, but projections of how future aquatic ecosystems will function are necessarily based on the traits of primary producers predicted to survive and even thrive in a high CO_2_ world. These projected winners are taxa that respond to CO_2_ enrichment by increasing their population growth rates, such as the cyanobacterium *Synechoccocus* (2), some nitrogen fixers (39), and some picoeukaryotes (40). Indeed, most phytoplankton functional groups have members with positive growth rate responses to CO_2_ enrichment (3, 11). Projections of the roles of these “winners” in future ecosystems is currently based on the indefinite maintenance of increased population growth rates in response to environmental amelioration. In contrast, we show that growth rate responses can reverse in ameliorated environments, and that traits underlying life-history and ecosystem function can shift substantially even in a constant, high-quality environment. Figure 4 shows examples of how evolutionary trait changes associated with Prodigal Son dynamics can affect trait calculations used in projections. These examples are illustrative only, but highlight that projections that rely on maintaining the plastic response (green) can differ those that incorporate trait reversion (blue). For population growth rate (4A), the effect of trait reversion on the average trait value is ~5%, and well within the intra-group variation used in projections of phytoplankton responses to increases in pCO_2_ (2). In contrast, the effect of trait reversion on biovolume production is ~30%. The magnitude of effects here are for a laboratory study and cannot be extrapolated to natural populations, but our results illustrate that trait reversion has the potential to dramatically alter projections of functional traits of future populations in ameliorated environments.

**Figure 4.**
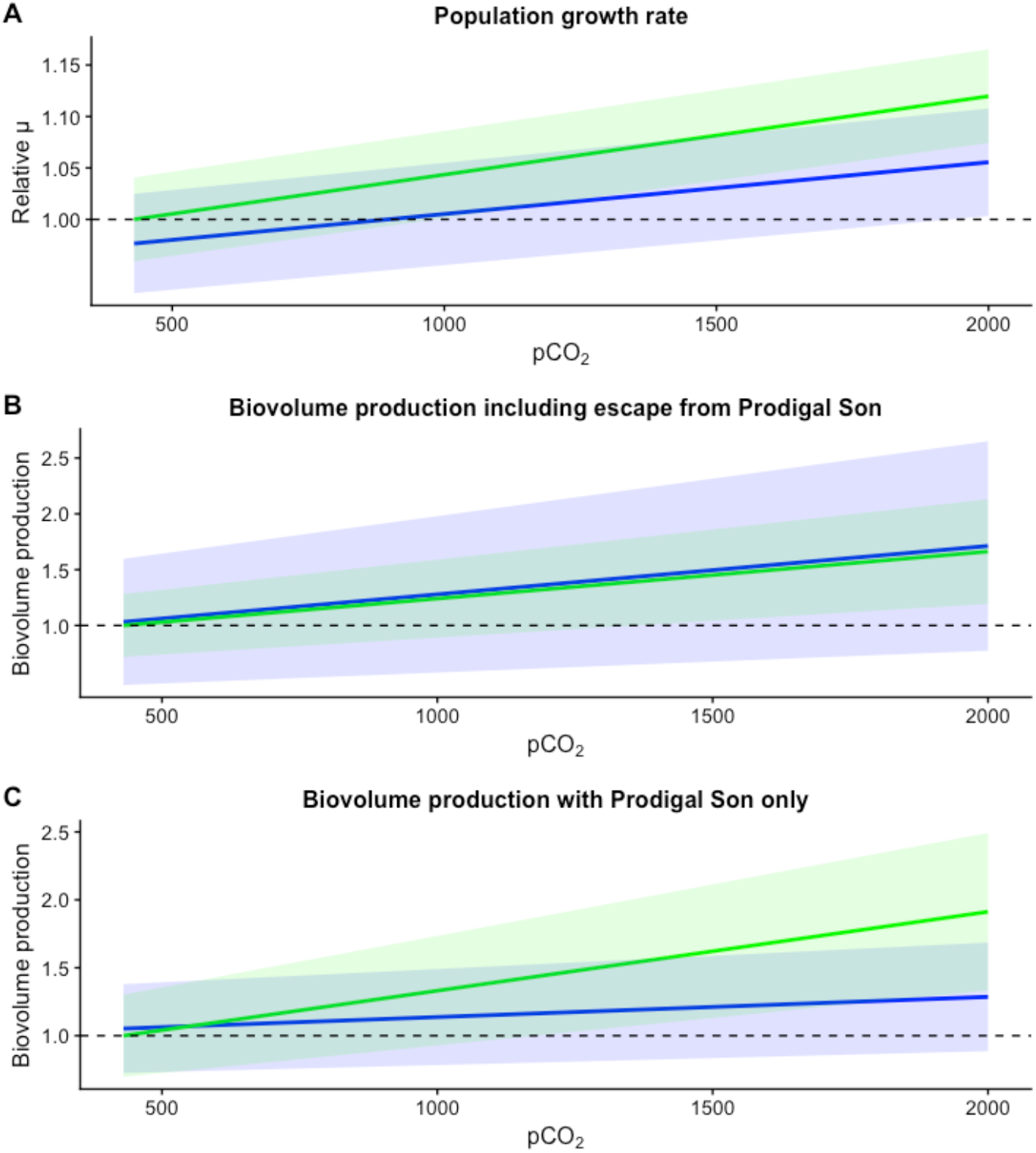
Visualization of statistical models of trait values, illustrating how projections can be affected by evolutionary responses. Lines indicate change in relative population growth (A) and relative biovolume production per unit time (B-C) explained by pCO_2_. Models fit on Amb populations are in green; models fit on High populations are in blue. Shading represents variation in model explained by genotype effects and is constant over pCO_2_ levels. Data are as in previous figures, details of statistical models are in SI. In all cases, the average trait value over all strains for the Amb populations at ambient pCO_2_ has been set to 1.0; other values are relative to this. In (A), evolutionary responses to high pCO_2_ reduce growth rate over all levels of pCO_2_, but slopes are similar. All five genotypes are included in this model. In (B) all five genotypes are also included in the model. Strain cc125 did not reduce fecundity or cell size, in contrast to other strains, after evolving under high pCO_2_. The evolutionary response to high pCO_2_ increases the variance in projections, but does not change the average response relative to projections based on plastic responses alone. In (C), genotype cc125 is excluded; all other genotypes show evidence of Prodigal Son dynamics. Here, evolutionary responses decrease biovolume projections relative to using plastic responses to changes in pCO_2_ alone.

### Conclusions

Projections of “climate change winners” are based on scenarios where “winners” currently exist in environments to which they are reasonably well adapted, and then experience environmental change as amelioration. Since projected “winners” are lineages or functional groups that increase in frequency, they must increase their relative growth rates; one way to do this is to increase absolute population growth rates. We show that when the environment goes from good to better and then remains improved over timescales long enough for evolution to occur, a tradeoff between producing high quality offspring and high numbers of offspring can manifest. In the case of this photosynthetic microbe, the decrease in offspring quality in the ameliorated environment is linked to the production and metabolism of ROS, which is a necessary byproduct of photosynthesis. Populations evolved in the ameliorated environment produce more robust but fewer offspring, which is consistent with selection for tolerating and minimizing damage associated with toxic metabolic byproducts such as ROS at the expense of allocating fixed C to building new cells. Since the underlying metabolic changes involve conserved processes (photosynthesis and ROS production), these dynamics are potentially general in phytoplankton growing in persistently ameliorated environments.

Assessing how Prodigal Son dynamics can impact projections of traits in “climate change winners” depends on the interplay between interlineage competition (genotype sorting) and intralineage evolution. In this study, cc125 escapes Prodigal Son dynamics, but is not fast-growing relative to the other genotypes. If its growth rate in single-genotype culture is indicative of its growth rate in competition with other genotypes, it would fail to increase in frequency, despite having escaped evolving lower lineage growth rates. If damage accumulation is slow, a genotype with a large plastic response to amelioration, such as cc1952, could outcompete other genotypes, but then experience a decrease in population size when selection against producing fragile offspring intensifies. On the other hand, if Prodigal Son dynamics are rapid, genotypes with the largest plastic growth responses could fail to overgrow competitors before having to reduce their lineage growth rates, leading to the maintenance of more diverse, slower-growing populations in ameliorated environments.

This study raises the question of which plastic trait changes will revert over time through the action of natural selection. Ecosystem functions such as primary production are modelled using trait values (41), so projections of functional traits in future environments requires predictions of which traits are likely to evolve towards a new value, and which traits are likely to revert over hundreds or thousands of generations. This study begins to answer this question. If the trends in our experiment generalize, then it is reasonable to hypothesize that average cell division rates that far exceed historical average cell division rates cannot be maintained indefinitely without affecting other functional traits through emergent tradeoffs. Our rationale is that this is because the pathways that clean up toxic metabolic byproducts such as ROS have probably not been selected to maintain consistently high activity levels, and we speculate that in most phytoplankton, the episodic nature of conditions favourable to growth (the right combination of light, temperature and nutrients) means that cellular repair could take place during periods of slow or arrested cell division. This would further decrease selection on efficient repair during periods of rapid cell division. In ameliorated environments, natural selection initially favours rapid population growth, it eventually shifts to favour lowering the accumulation of and sensitivity to toxic metabolic byproducts, such as the reactive oxygen generated by photosynthesis, at the expense of allocating fixed carbon to growth. While the strategies used to accomplish this reduction in damage and ROS sensitivity vary between genotypes, some decrease in gross photosynthesis, usually coupled to a decrease in fecundity and population growth rate, is a general feature of the overall strategy for producing higher quality daughter cells in enriched environments. Thus, we hypothesize that trait reversion is more likely for metabolic traits involved in “dirty work” (13) – metabolic work that generates toxic byproducts. In particular, we hypothesize that dirty work pathways that speed up substantially in response to environmental amelioration are more likely to revert than are other traits that respond plastically to the same environmental change.

This work provides both evolutionary and physiological mechanisms for understanding how photosynthetic unicells respond to environmental amelioration. More broadly, it sheds light into selection pressures that act during sustained, rapid growth in phytoplankton, such as algal blooms. Taken together with observations that the production of the toxin microcystin during algal blooms (42) may function to stabilize proteins during the late stages of algal growth, this study points towards a general role of natural selection acting to mitigate the effects of toxic metabolic byproducts on cells during rapid population growth. Here, instead of producing compounds that mitigate the effects of toxic metabolic byproducts on proteins, our populations evolved the ability to tolerate and metabolise ROS. Understanding how tradeoffs between functional traits emerge during sustained rapid population growth in phytoplankton opens up a new perspective on understanding how and why bloom-forming algae evolve, as well as how natural selection acts on “climate change winners” over hundreds of generations.

## Methods summary

Detailed methods, calculations, and statistical models are in the supplementary material.

### Culturing methods and selection experiment

All populations were started from single cells of *Chlamydomonas reinhardtii* provided by N.Colegrave (University of Edinburgh). Genotypes are catalogued as cc125, cc1691, cc1952 and cc2937 in the Chlamydomonas Resource Collection. Six independent replicate populations of each genotype in each environment were grown, for a total of 60 populations in the experiment. All populations were grown in sterile microwell plates in 2mL of Tris-Acetate-Phosphate (TAP) media (43). pCO_2_ levels in media were manipulated by pre-bubbling media with air of the desired pCO_2_ and then keeping growing plates in CO_2_ controlled incubators (15). Every two days the 1% of the each population was transferred to a fresh well with pre-bubbled TAP media. All populations were grown at 25°C under continuous light at 170 μmol photons/sec/m^2^ and shaking at 100rpm. The entire selection experiment lasted 90 transfers, which corresponds to approximately 900 generations in the High populations.

### Plastic and evolutionary trait responses

For all trait assays, populations were acclimated to the assay environment for one growth cycle. For all traits measured, the plastic response is calculated by comparing the trait values of a given population in ambient and high pCO_2_. For example, the plastic growth response of a population, regardless of the regime where it evolved, would be calculated by comparing growth at high pCO_2_ and growth at ambient pCO_2_. The evolutionary response is calculated by trait values of populations evolved in the high pCO_2_ environment with the trait values of the same genotype evolved in the ambient pCO_2_ environment when grown in a common environment. For example, the evolutionary growth response to high pCO_2_ is calculated using the growth rates of High and Amb populations of the same genotype in the high pCO_2_ environment.

### Population growth rates

For all trait assays, populations were acclimated to the assay environment for one growth cycle. For population growth rate measurements, populations were diluted after acclimate in the assay environment and an inoculum of ~40 000 cells were transferred to fresh media at the assay pCO_2_. Population growth was calculated after 48 hours as ln(N_48h_/N_0h_)(44). Cell number was quantified by measuring absorbance at 750nm and converting to cell number using genotype and environment-specific standard curves.

### Cell fecundity

*C. reinhardtii* divides by multiple rounds of binary fission(45). The number of daughter cells per mother cell was measured by fixing cells on agar and incubating them in the dark, which arrests cell growth and initiates binary fission (29). Populations were grown to midexponential phase in the relevant environments, diluted 10-fold, and plated onto 1.5% TAP agar plates. The plates were incubated in the dark for 12 hours at 25°C. The number of daughter cells per mother cell was scored under a microscope. Time needed for a single cell cycle was calculated from population growth rates and average fecundity.

### ROS tolerance

ROS tolerance was assayed on a gradient of Rose Bengal (RB) ranging from 0 to 4.5μM. Growing populations were inoculated into fresh growth wells containing TAP and RB and growth was measured over one growth cycle. RB resistance as the amount of RB needed to reduce population growth by 50% was then calculated using a sigmoid decay function.

### Other trait measurements

Cells were counted and cell size was measured by flow cytometry using size-bead calibrated forward scatter. Photosynthesis and respiration rates were measured for each population using 2-mL oxygen optodes (PreSens) in pre-bubbled media at the relevant pCO_2_ level.

### ROS gene expression

Gene expression for *Cat*, *Gpx* and *Fsd1* were measured using quantitative PCR (qPCR) using previously developed primers(34–36). Primer information is in SI. RNA from independent triplicate acclimated cultures for each population was extracted as per(46). cDNA was synthesized using Bioline’s Tetro cDNA kit and cDNA libraries were stored at −20C°. qPCR reactions were prepared using a GoTaq qPRC kit (Promega).

### Statistical analyses

All statistical analyses, including the models used to produce Figure 4, are detailed in the SI and were carried out in the R environment (final analyses carried out in version 3.5.0, R Core Team (2018). All code will be made available at the time of manuscript acceptance.

## Supplementary Information and detailed methods

### Cell fecundity measurements

*C. reinhardtii* divides by multiple binary fission(1). The number of daughter cells per mother cell was measured by fixing cells on agar and incubating them in the dark, which arrests cell growth and initiates binary fission(2). Populations were grown to midexponential phase in the relevant environments, diluted 10-fold, and plated onto 1.5% TAP agar plates. To control for differences in the number of cell divisions that could occur in 12 hours, cells were size-selected by centrifugation at 600rpm for 20 minutes(2, 3). Larger cells are spun to the bottom of the tube while the top layer of media contains smaller cells. We tested the effectiveness of size selection by comparing the distributions of cell division numbers on plates with and without size selection, and found that size selection was effective; plates containing large cells showed evidence of colonies that had gone through >1 cell cycle, while colonies on size-selected plates were made up of protoplasts inside unruptured cell walls. Plates were incubated in the dark for 12 hours at 25°C. The number of daughter cells per mother cell was scored under a microscope at 400x magnification. Note that is it possible for daughter protoplasts to undergo fission at different rates, so that the numbers of protoplasts inside a mother cell is not always 2^n^. For example, if the mother cell divides, and then the daughter protoplasts divide at different speeds, it is possible to observe 3 protoplasts inside the cell wall.

### Cell size

Cells were measured using flow cytometry (BD LSRFortessa). Forward scatter area was equated to cell diameter by a standard curve using microbeads. The parameters of the curve are Diameter μM = x9.01x 10^5^ + 0.28873 where x is the value for forward scatter. Average cell diameter per population was based on a sample of at least 10,000 cells. Volume was calculated assuming cells are spheres.

### Photosynthetic oxygen evolution and consumption

Photosynthesis and respiration rates were measured for each population using 2-mL mini sensors (SensorVial SV-PSt5-2mL, Sensor dish reader SDR2, PreSens). O_2_ levels were recorded using optical fluorescence-based oxygen respirometry. Each culture was measured at mid-exponential phase in 1x volume of fresh TAP bubbled with the appropriate CO_2_ level in a 2 mL glass sensor vial. The vials were filled to the brim and sealed with cling film, thus preventing gas exchange with the atmosphere or air bubbles in the vial. All samples were first incubated in the dark for 20 minutes. The vials were then placed on the 24-channel oxygen meter and O_2_ evolution measured for 3 minutes every 15 seconds on a shaking table. The vials were then mixed by inversion and immediately wrapped in tin foil. O_2_ consumption was measured for 3 minutes every 15 seconds. The sensor dish reader was connected to the PreSens data logger and data collected in the SDR_v4 software. Data were then processed in R to calculate oxygen evolution and consumption rates by fitting slopes to oxygen levels during the appropriate time windows. Rates are μmol O_2_ per hour per cell. Oxygen evolution and consumption rates are normalized to cell number. Following measurements in the mini sensors, 200μl of culture was removed for cell counting. Cells were counted using flow cytometry on a BD LSRFortessa.

### ROS resistance

Resistance to ROS was assayed on a gradient of Rose Bengal (RB) ranging from 0 to 4.5 μM. In its liquid form RB is split into the oxygen radical O-when excited by light. We calculated the amount of RB required to half population growth in liquid culture, *xmid*. The cultures were first acclimated to assay CO_2_ environment for eight generations and then diluted to the same density and aliquoted into the RB gradient. *xmid* was calculated from OD 750 nm after 24 hours of growth using a logistic decay function:

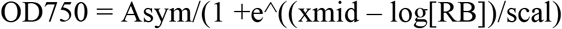

Where Asym is the maximum cell density, [RB] is the RB concentration, and scal a rate constant. Gradient assays were repeated thrice for each population and environment.

### Heat shock recovery

Populations were grown to mid-exponential phase and then diluted to 100 cells/mL. These are “initial” cells. 85μL per population was then placed in a PCR plate and incubated for 20 minutes at 44°C, followed by 10 minutes cooling at 25°C. Heat-shocked samples were then inoculated into standard growth wells and conditions and grown for one growth cycle, at which point population growth rate was calculated. Heat shock tolerance measurements were carried out in triplicate for each population in each environment reported. Heat shock recovery is given as the number of cells produced per initial cell per day for the 48 hours following the heat shock. See SI Figure 1.

### RNA extraction

RNA was extracted for populations grown at high CO_2_ only, so that comparisons are between Amb populations and High populations. Populations were acclimated to the high CO_2_ environment for two growth cycles. Three replicate cultures of each population in each environment measured were founded and distributed randomly onto 48-well plates. They were incubated at high pCO_2_ for 36 hours and then harvested for RNA extraction.

RNA was extracted following Lake and Willows (2003) (4) who have optimised RNA yield in *C. reinhardtii*. Populations were harvested by centrifugation at 6000 rpm for 15 minutes. The centrifuged growth medium was discarded and the pellet immediately resuspended in RNAlater. Samples were stored at 4 °C until RNA extraction. To extract RNA, samples were pelleted at 13000 rpm for 15 minutes and RNAlater discarded. The pellets were resuspended in 0.5 ml of extraction buffer (50 mM Tris-HCl pH 8.0, 0.3 M NaCl, 5 mM EDTA and 2% w/v SDS, proteinase K 40 μg/mL) and incubated on an orbital shaker at room temperature for 20 min. To extract total RNA, an equal volume of phenol:chloroform (125:24:1) (phenol chloroform equilibrated to pH 4.3) was added, vortexed, and centrifuged at 2000 rpm for 5 min. Both the bottom layer, containing proteins and cellular debris, and the DNA-containing cloudy interface were discarded. The upper aqueous phase, containing the RNA, was re-extracted with phenol:chloroform four times, or until there was no cloudy interface left in the extract. The RNA was precipitated in 2 volumes of 100% ethanol at −20°C overnight and recovered by centrifugation at 8500 rpm at 4 °C for 15 min. Pellets were washed with 70% ethanol, centrifuged for a further 10 min and air dried at room temperature for 1 hour. Pellets were resuspended in RNase free water and stored at −80 °C.

### cDNA preparation and quantitative PCR (qPCR)

Complementary DNA was synthesised using Bioline’s Tetro cDNA synthesis kit. Five μL of RNA extract was mixed with 1 μL of Oligo dT Primers, 0.5 mM dNTPs, 1X RT buffer, 1 uL of RiboSafe inhibitor and 10 units of Tetro Reverse Transcriptase, final concentration. DEPS-water was added to 20 μL. The reaction was prepared on ice, vortexed, and incubated for 30 minutes at 45 °C. The reaction was terminated by further 5 minutes of incubation at 85 °C. cDNA libraries were stored at – 20°C until used. qPCRs were done in StepOnePlus thermocycler with Promega’s GoTaq qPCR kit. The reaction volume was 20 μL with the following ingredients: 0.9 mM of forward and reverse primers, 1X GoTaq qPCR mastermix, 0.2 % v/v CRX reference dye, 2 μL of cDNA template, and nuclease free water to 20 μL. The amplification protocol was 10 minutes at 95°C, then 40 cycles of 95°C for 15 seconds, 65°C for 30 seconds and 72°C for 30 seconds. Target antioxidant gene expression products and their primers are listed in SI Table 1.

All PCR quantification and statistical analyses were carried out in R(5). Fluorescence threshold Cq was determined by the second derivative of a five parameter sigmoid curve using the qPCR package(6). This value was estimated for each technical replicate separately. The efficiency of each amplification was estimated at the maximum of the second derivative.

Values shown are relative cDNA per cell, calculated as (average High-L)/(average Amb-L) for each independently evolved population. Averages are made using independent growth and RNA extraction events for each evolved population. Error propagation was calculated using simple relative errors, as errors was very small relative to average values.

### Statistical analyses

All statistical analysis and the construction of figure 4 were carried out in the R environment(5). All R code is available upon request and will be deposited at the time of manuscript acceptance. The nlme package was used for the analyses below.

All statistical models for traits other than population growth were mixed models, with assay CO_2_ level (high or ambient) and selection history (High or Amb) as fixed effects, and genotype as a random effect (random slope only, due to the low number of genotypes used in this study). For population growth, the number of daughter cells produced per parent cell per cell cycle (“daughters”) was also included as a fixed effect. Interactions between fixed effects were included in all models.

To determine whether the “daughters” term should be included in the model for population growth rate, we first constructed a full model with assay CO_2_, selection history, and daughters as fixed effects, with interactions, then removed non-significant interactions. We then compared the models with and without daughter included were included using Akaike information criterion.

| Details | Model number | df | AIC | p-value |
| --- | --- | --- | --- | --- |
| **Daughters** | 1 | 10 | **-787.19** |  |
| No Daughters | 2 | 7 | -776.56 | 0.0008 |

For the conceptual Figure 4, separate models were constructed for the High and Amb populations with population growth (μ) as the response variable. In each case CO_2_ level was the fixed effect and strain was a random effect. The visualization used the slope of the fixed effect as slope, and the sd of the random effect as the sd on the plot. In all cases, the values were normalized so that the values used to contruct visulalization the Amb scenario (green) had an intercept of 1.0, so that the figure shows fold changes, rather than absolute values. In panel C, the statistical model was run with cc125 excluded, but the method was otherwise identical to panel B.

**Supplementary Figure 1.**
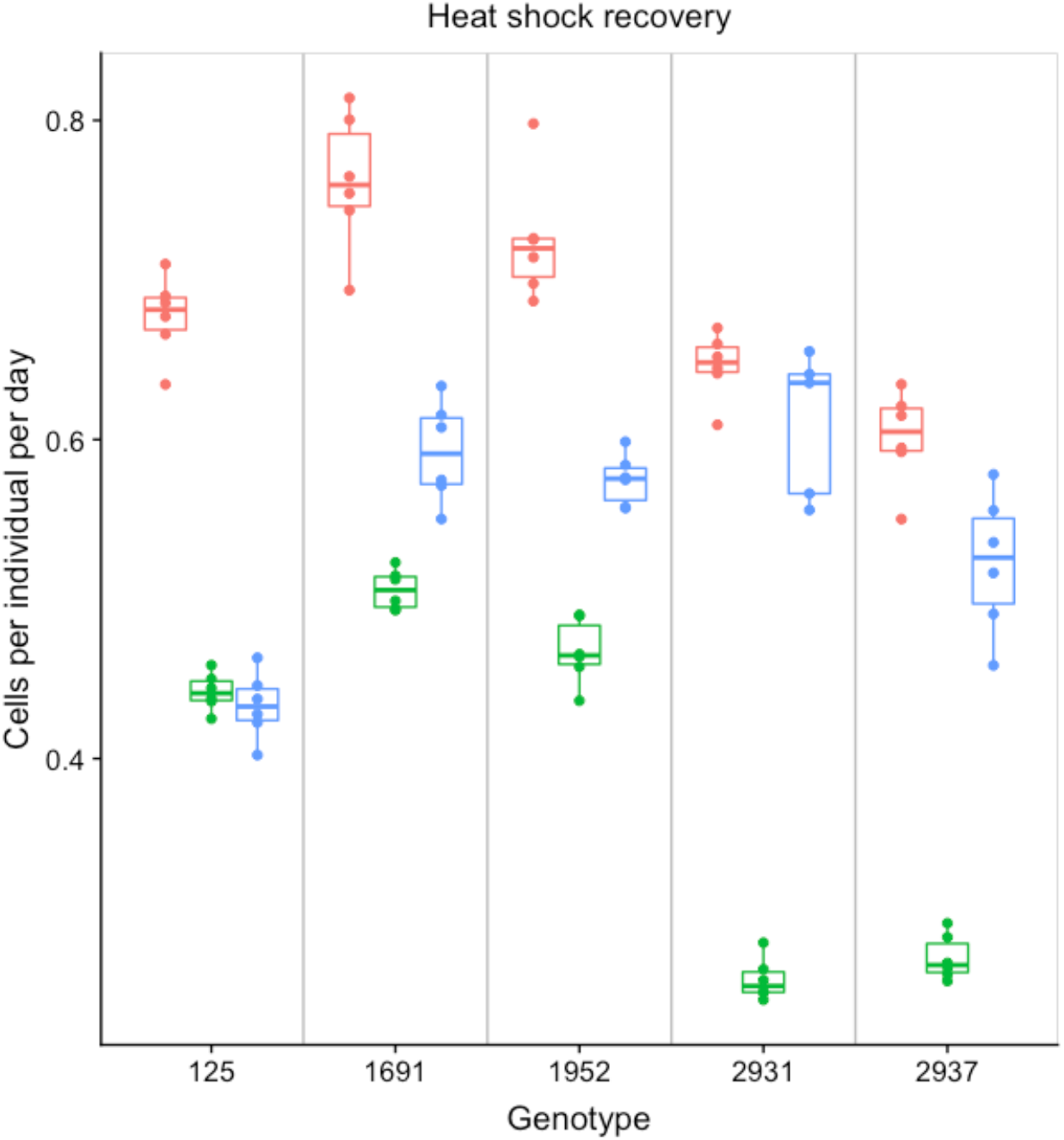
Daughter cells produced per initial cell per day following heat shock. Trait values in pink correspond to the Amb-L populations growing in ambient CO_2_, and correspond to the baseline trait prior to environmental amelioration. Trait values in green show Amb-L populations growing in high CO_2_, which corresponds to the plastic response to environmental amelioration. Trait values in blue show High-L populations growing in high CO_2_, and are the evolved phenotype after ~900 generations in the ameliorated environment. Each point represents the average trait value for a single replicate population, boxes show means for each genotype. In general, Amb-L populations grown in ambient pCO_2_ recover best from heat shock (pink values). Short-term exposure to high pCO_2_ decreases heat shock recovery (green) relative to the baseline values in pink. However, after ~900 generations in the high pCO_2_ environment, heat shock recovery increases (blue). In contrast to ROS tolerance in Figure 1H where short term exposure to high pCO_2_ is protective, here is it detrimental. This is consistent with ROS metabolism being stimulated, but cells being in worse overall condition, even after only ~9 generations in the high pCO_2_ environment.

**Supplementary Table 1.**
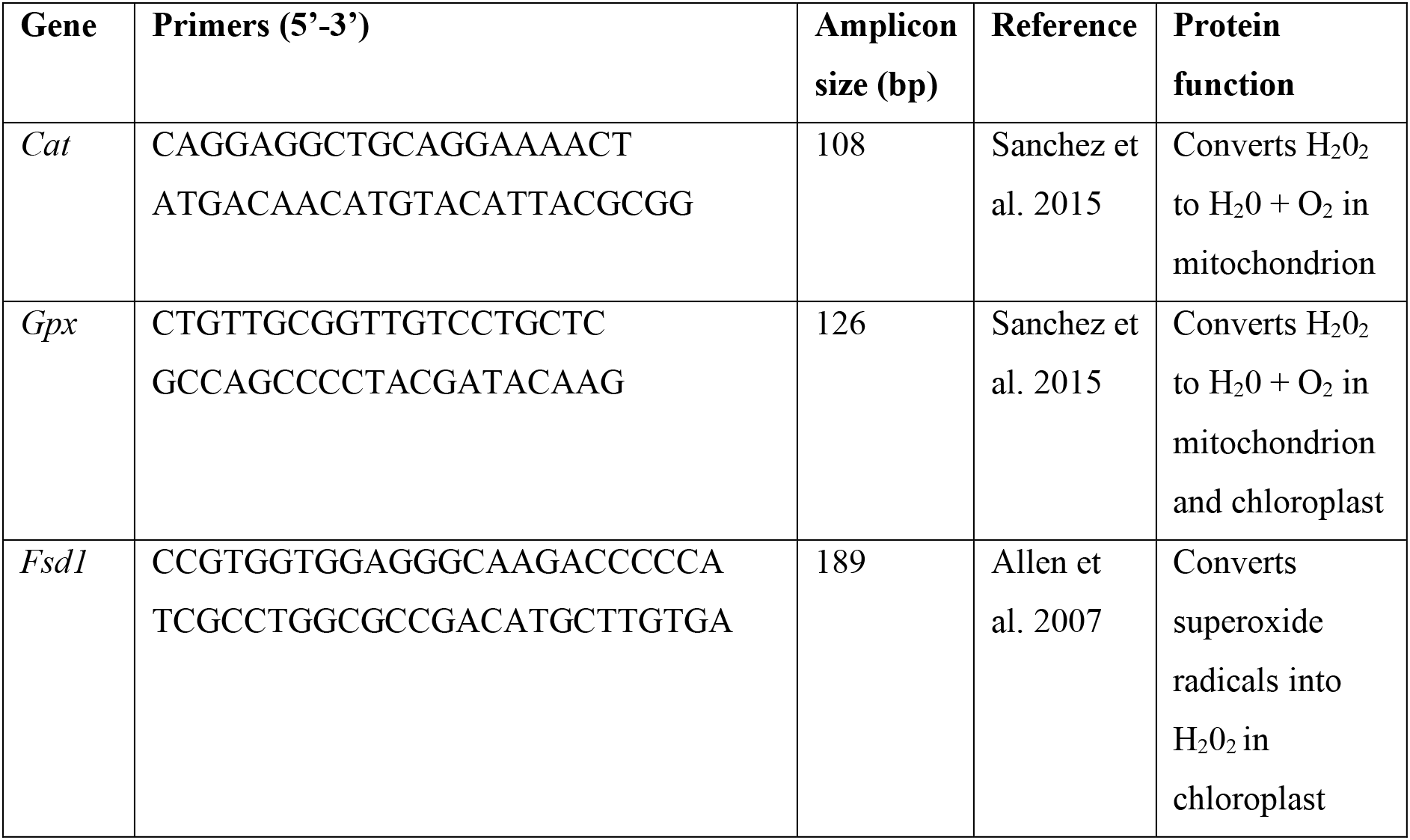
Primers used for qPCR based measures of ROS metabolism gene expression levels.

